# Variant Calling Parallelization on Processor-in-Memory Architecture

**DOI:** 10.1101/2020.11.03.366237

**Authors:** Dominique Lavenier, Remy Cimadomo, Romaric Jodin

## Abstract

In this paper, we introduce a new combination of software and hardware PIM (Process-in-Memory) architecture to accelerate the variant calling genomic process. PIM translates into bringing data intensive calculations directly where the data is: within the DRAM, enhanced with thousands of processing units. The energy consumption, in large part due to data movement, is significantly lowered at a marginal additional hardware cost. Such design allows an unprecedented level of parallelism to process billions of short reads. Experiments on real PIM devices developed by the UPMEM company show significant speed-up compared to pure software implementation. The PIM solution also compared nicely to FPGA or GPU based acceleration bringing similar to twice the processing speed but most importantly being 5 to 8 times cheaper to deploy with up to 6 times less power consumption.

## I. Introduction

With an estimated 100M of human genomes to be sequenced in 2025, the computing challenge to process the Terabytes of data produced by sequencers is at the root of the upcoming revolution in personalized medicine. Both the speed and the cost of genomics analysis will be determinant in the wide spreading of its applications.

A fundamental genomic analysis, called *variant calling*, consists in detecting, at a DNA level, small differences between two genomes. More precisely, from a pool of short DNA fragments (reads) coming from a specific individual, and obtained using high throughput sequencing, the objective is to locate the place, in a reference genome, where short DNA strings (< 50 bp) differ. The variant calling process proceeds in several steps which first map the reads on the reference genome and then analyse the mapping especially in the regions where differences are found.

Many software, such as GATK [1], Strelka2 [2], Varscan2 [3], SOAPsnp [4] or DeepVariant [5] have been developed for that purpose. Although these tools have some advantages and disadvantages, they are daily used to identify a large number of specific variations in many health or agronomic application domaines. Due to the large volume of data to analyze, the execution times of these software can be very long, i.e. a few hours on standard bioinformatics servers. Thus, speeding-up the variant calling process is a real challenge, especially in the context of personalized medicine that requires systematic deep analysis of human individual genomes.

Several methods have been proposed to accelerate variant calling by the means of parallel and distributed computing techniques: HugeSeq [6], MegaSeq [7], Churchill [8] and Halvade [9] support variant calling pipelines related to GATK [10]. These parallel implementations exploit the fact that the alignment of one read is independent of the alignment of the others, while the call of variants is independent from one region to another. Other parallel pipelines for variant calling include SpeedSeq [11] and ADAM [12].

Other strategies are based on hardware accelerators. FPGA technologies are particularly well suited to hardwire DNA computation intensive algorithms such as sequence alignments or read mapping. Among recent FPGA systems dedicated to genomic data analysis, the following platforms demonstrate significant speed-up compared to standard GATK software: the Illumina DRAGEN-Bio-IT platform [13] and the WASAI Lightning platform [14]. These FPGA architectures associate both reconfigurable computing resources and memory chips. They provide nice speed-up ranging from 10 to 50 on variant calling applications.

GPU devices offer another alternative to reduce genomic analysis runtime, especially for read mapping which is an important step in the variant calling process [15][16][17][18]. More recently the parallelization of GATK on the NVIDIA Clara Parabricks pipelines [19] achieves a 35-50X acceleration.

This paper explores another way of speeding up the variant calling process using a Processing-in-Memory (PIM) architecture. We present an original parallelization based on new PIM chips developed by the UPMEM company. Actually, PIM architecture is not a new concept. In the past, various research projects have investigated the possibilities to close data and computation. The Berkley IRAM project [20] probably pioneers this kind of architecture to limit the Von Neumann bottleneck between the memory and the CPU. The PIM project of the University of Notre Dame [21] was also an attempt to solve this problem by combining processors and memories on a single chip.

UPMEM solution tackles the problem by designing high density DRAM and RISC processors on the same die. Several chips are then encapsulated into standard 16 GBytes DIMM modules. The idea is to complement the main memory of a multicore processor with PIM devices. Data located in these specific memories can be independently processed releasing the pressure on the CPU-memory transactions.

The variant calling task perfectly illustrates how such time-consuming applications can benefit from the PIM architecture. The mapping step, which represents a large part of the overall computation time, is particularly well suited, as fine grained parallelization can be efficiently executed to perform multiple independent alignments along the whole reference genome. Deporting this activity directly to the PIM-DRAM module, and parallelizing the whole process to hundreds of PIM cores, avoids a lot of CPU-memory transactions compared to a standard multithreaded solution.

The objective of the research work presented here, is to precisely evaluate the potentialities of a PIM architecture composed of a bunch of UPMEM DIMM modules, coupled to the main computer memory bus, on a critical genomic treatment. A generic variant calling algorithm, called upVC, has been implemented as a testbed on real PIM components to provide exact measurement and fair comparison with existing systems in terms of speed-up, energy consumption and cost. From a quality point of view, the upVC implementation is not intended to immediately compete with mature software such as GATK.

The rest of the paper is structured as follows: the next section describes the main features of the PIM architecture proposed by the UPMEM company. Section 3 details how the variant calling process is implemented on a PIM architecture. Section 4 details the experimentation conducted on a 1024 core prototype PIM. Section 5 compares the PIM approach with alternative hardware accelerators. Section 6 concludes the paper.

## II. UPMEM Architecture Overview

### A. Server level architecture

UPMEM’s PIM technology consists of thousands of parallel coprocessors (called DPU) within the main memory of a host CPU (e.g., x86, ARM64, or Power9). Standard and UPMEM DIMMS can coexist on a server to operate both regular processing and PIM. The CPU provides programming instructions to DPUs, and collects their results as they operate individually. This design relieves the CPU from a memory bottleneck and greatly reduces the energy hungry data movement.

**Figure 1:**
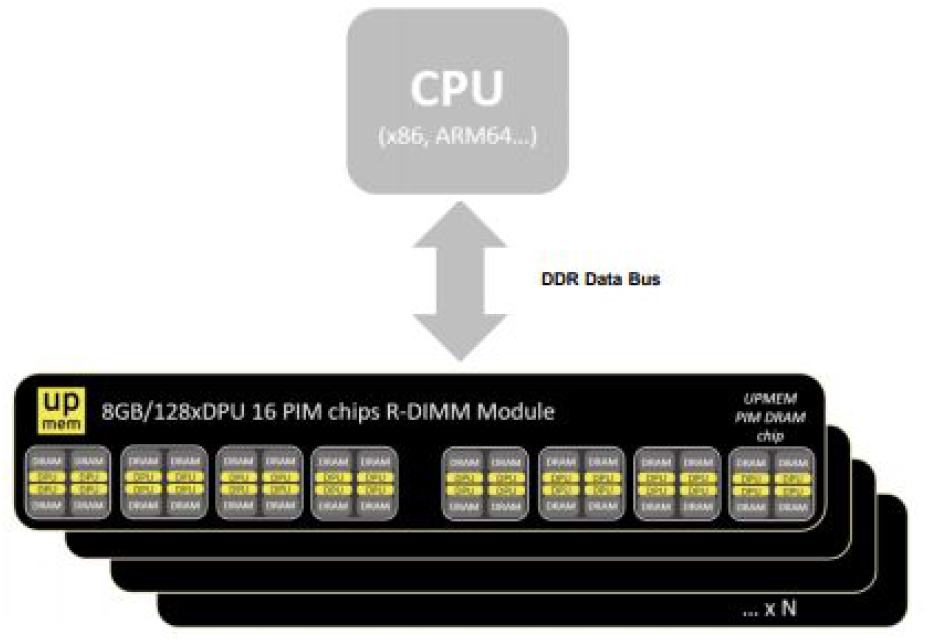
PIM server architecture

### B. DIMM organization

The UPMEM DDR4 2400 DIMM (dual ranks) comprises 16 PIM enabled chips totaling 128 DPU. A DPU is a 32-bit processor running at 500MHz. Up to 20 UPMEM DIMMs can be plugged into a x86 platform, keeping 2 slots per socket for traditional DRAM. The solution scales with the ability to increase the number of DPU in a system and can reach 5120 DPUs in a quadri socket platform with 40 PIM DIMMs.

### C. The chip

A PIM memory chip contains 8 DPUs. Each DPU is associated with 64MB of DRAM shared with the host CPU. The calculations happen on chip and within each unit with a memory bandwidth of 1GB/s. This is why PIM imposes data locality and parallelism with the consequence to alleviate the need for important data movements.

### D. The DPU

A DPU is a 24 threads, 32-bit RISC processor – with 64-bit capabilities – working at 500Mhz with an ISA close to traditional ARM or RISC-V equivalent processors, making it easily programmable. DPUs have a 64KB of WRAM (Working RAM) and a 24KB instruction memory, called IRAM, that can hold up to 4,096 48-bit encoded instructions. DPUs are independent from each other and run asynchronously

**Figure 2:**
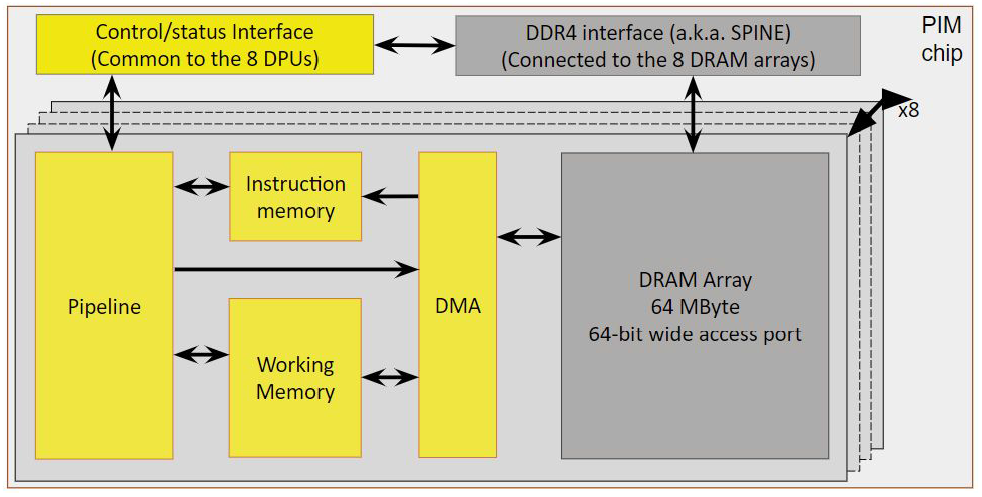
UPMEM PIM chip diagram

### E. Development Environment

Every DPU can be programmed individually or in groups while orchestrated through the host code. The PIM architecture sits on an efficient toolchain centered around a LLVM based C-compiler using LLVM v10.0.0 and with Linux drivers for x86 servers. It also contains a full-featured runtime library for the DPU, management and communication libraries for host to DPUs operations and a LLDB based debugger. This experiment has been achieved using the SDK v2020.3.0 [27].

## III. Variant calling on PIM

This section presents the variant calling strategy elaborated to fully exploit the PIM architecture. The next subsection gives first an overview of the implementation, and how the full variant calling process has been parallelized.

### A. Overview

Schematically, the variant calling computation is based on a main loop that sequentially process packets of reads:

~~~
   Loop
      G: Get paired-end read packet from disk
      D: Dispatch read packet to DPUs
      M: Map reads on DPUs
      U: Update VC data structure
~~~

The loop is composed of 4 independent tasks. The first one (G) get packets of paired-end reads from external storage support, the second one (D) dispatched these reads to the DPUs, the third one (M) maps the reads using the DPU computational resources to the reference genome, and the last one (U) updates a variant calling (VC) data structure with the mapping results of the previous stage.

Parallelism is both brought by pipelining the four tasks of the loop on the main processor and by executing concurrently the read mapping on the PIM memory. The 4 tasks overlap as follows:

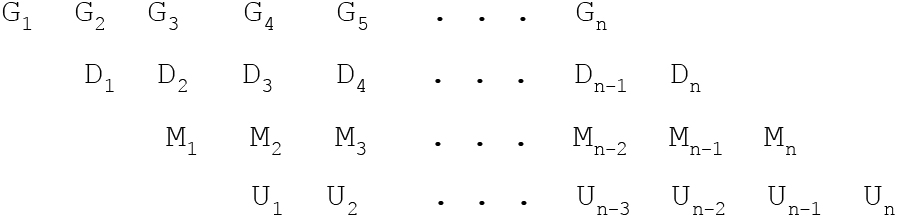

Considering *n* packets of reads, the total loop execution time is approximately equal to t_max_ x (n+3), where t_max_ is the time of the slowest task. In this implementation, a first challenge is to have a good load balancing between these tasks in order to fully exploit the parallelism capacity of the multicore processor together with the PIM memory computing resources.

Before entering the loop, the reference genome is indexed based on small kmers (see genome indexing section). The index is then distributed non redundantly across all the DPUs. This initialization step (genome indexing + index dispatching) can be significantly long, compared to the loop execution. However, in the case of intensive human variant calling, for example, it can be performed only once. Different sequencing datasets can then be processed without resetting the PIM memory.

The final variant calling stage is performed after the loop execution. The U task of the loop simply updates a specific data structure according to the results of the mapping alignments. This is a fast process that consists in scanning intermediate results built during the U task.

### B. Genome indexing

As the mapping is done by the DPUs, the idea is to build a distributed index allowing reads to be evenly dispatched into the 64 MB memories associated with each DPU. The index is a hash table whose *keys* are all the distinct kmers of the reference genome and *values* are a list of positions of the kmers (chromosome number, position on the chromosome) associated with their nucleotide context.

|  | substitution |  | insertion |  | deletion |  |
| --- | --- | --- | --- | --- | --- | --- |
|  | upVC | GATK | upVC | GATK | upVC | GATK |
| TP % | 99.77 | 100 | 99.10 | 99.69 | 99.57 | 99.98 |
| FN % | 0.50 | 0.23 | 0.91 | 0.74 | 0.57 | 0.37 |
| FP % | 0.23 | 0 | 0.90 | 0.31 | 0.43 | 0.02 |
| P % | 99.77 | 100 | 99.10 | 99.69 | 99.57 | 99.98 |
| S % | 99.50 | 99.77 | 99.08 | 99.26 | 99.42 | 99.63 |

The nucleotide context corresponds to the nucleotide sequence right after the key sequence on the reference genome. Its size is equal to the read size.

kmer size has been fixed to 14, leading to 268,435,456 (*4^14^*) different entries in the hash table. These entries are dispatched in the different DPUs in such a way that each DPU memory will be allocated with the same amount of data. As the number of DPUs is smaller than the number of entries, each DPU will contain several entries. The entry assignment is done according to a workload estimation based on the genome kmer distribution.

### C. Mapping

After receiving a read packet, reads are dispatched to the DPUs. As the first 14-mers of the reads correspond to the key, it can be immediately sent to the DPU where the corresponding entry has been allocated. This is the D task of the loop. Once all reads have been dispatched, each DPU contains a subset of reads that are ready to be mapped.

The mapping algorithm proceeds in two stages. First a no gap alignment algorithm is performed. This is a very fast procedure that finds good maps with the majority of the reads. It simply computes a Hamming distance between a read and its associated nucleotide contexts. For example, the read AAAATTGGAGCTACAGCGT whose 4-mer key is `AAAA`, will trigger Hamming distance computation between the two sequences corresponding to the first entry of the hash table of figure 3.1.

**Figure 3:**
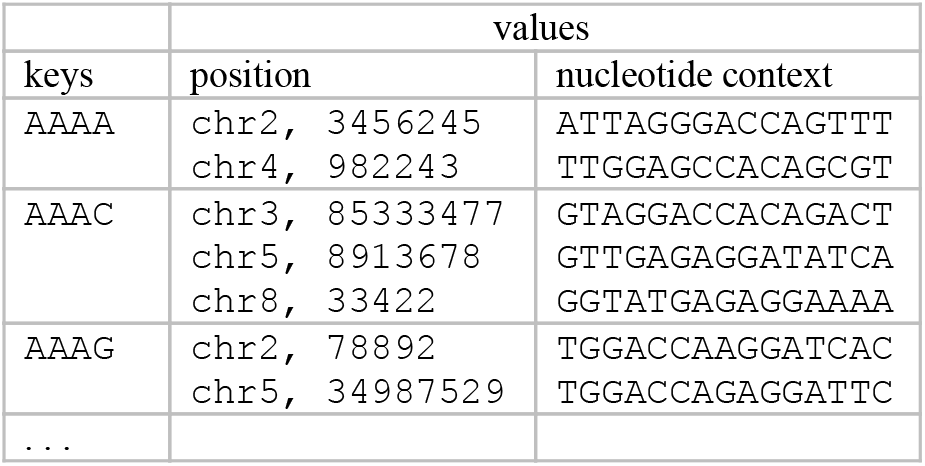
hash table example

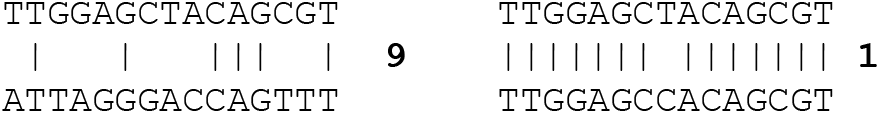

The procedure is optimized in such a way that the computation of the Hamming distance is stopped when it goes beyond a predefined threshold. Furthermore, at that point, we search for potential indels by shifting from 1 to 5 positions the 2 sequences, and by comparing the next 4 nucleotides. If there is a match, then the second stage is triggered.

The second stage executes a banded Smith & Waterman algorithm that precisely locates indels between the two sequences. The result of the mapping step is the position of the reads where good maps have been found together with their scores. Several mapping locations can be attached to a single read.

### D. Variant Calling

The variant calling is done once all the reads have been mapped. However, during the loop execution, the U task preprocess the mapping results of the previous packet of reads. First of all, according to the mapping position of a pair of reads, it finds the best location of the pair on the reference genome. If several locations map identically, then the position with the lowest coverage on the genome is chosen to ensure a regular distribution of the read mapping.

In addition, the U task performs the following actions:

- **Update the coverage**. Each chromosome is associated with an integer array of the same size. This array is zero initialized. Each time a read maps the chromosome, the array entries corresponding to the mapping positions are incremented by one. For example, if a read maps at position 246 on chromosome 3, then the entries [246:246+read_size] of the array associated to chromosome 3 are incremented by one.
- **Update a list of variants**. Each read is systematically aligned to the reference genome at the position previously chosen. The alignment provides potential variant positions which are systematically stored in a list. An element of the list is defined by the 3-uplet *<position, type, occurrence>.* If a new variant is detected, a new element of the list is created. If not, the occurrence item is incremented by one.

After the full loop execution, the list of variants is scanned and decisions, based on the *occurrence* item, are taken to consider if elements of the list are effective variants, or sequencing errors.

Performing variant calling task in parallel with the mapping task has the following advantages:

- The major part of the variant calling computation time is hidden. Only the last part (scan of the variant list) is visible. Actually, it represents a very small fraction of the time dedicated to the variant calling computation.
- No intermediate file, such as BAM file, for storing the mapping results is needed. Sorting the alignments is also no longer needed as it is done in conventional calling variant pipelines where the mapping step and the variant calling step are done sequentially.

### E. Implementation

The parallelization described on the previous section supposes the reference genome index to fit completely into the PIM memory. If the number of available DPUs is too small, then the variant calling process must be partitioned into several passes. The current implementation includes this possibility. The structure of the program becomes:

~~~
   Loop (external)
      I: load part of index into DPU
      Loop (internal)
          G: Get paired-end read packet from disk
          D: Dispatch read packet to DPUs
          M: Map reads on DPUs
          S: Store map results
          U: if last external loop
              Update VC data structure
~~~

In this configuration, the index is split into P parts and the external loop is run P times. The full read dataset is then processed P times on different parts of the index This implies to slightly modify the update of the VC structure which can only be done after all reads have faced the full index. A new task (S) is thus added to store intermediate mapping results, and the VC update is simply done on the last iteration. The length of the internal loop pipeline is now equal to 5.

Another level of parallelism is achieved by multithreading the tasks D, S and U. Their parallelization with 8 threads ensures a short time execution for each of them.

This optimized implementation is written in C and is called upVC in the rest of this paper.

## IV. Experiment with a 1024 DPU System

This section describes the experiment conducted on a real 1024 DPU prototype system (266 MHz) for calling variants on the Human genome using the upVC implementation.

### A. Dataset

In order to have a ground truth to validate the correct functionality of upVC, and also to compare results with other variant calling softwares, a simulated sequencing dataset has been generated according to the following steps:

- **step_1**: The DNA sequence has been extracted from the HG38 Human reference genome^1^.
- **step 2**: A list of 3,153,377 variants from the common_all_20170710.vcf file^2^ of the dbSNP database has been created by randomly selecting variants indexed in this file. Variants have been restricted to SNPs and small indels (<= 5nt).
- **step_3:** Paternal and maternal chromosomes have been generated using the vcf2diploid tool [22]. Input data are the human genome reference sequence (step_1) and the list of variants selected at step_2.
- **Step_4:** Short Illumina paired-end reads have been generated with the ART read simulator [23] for all chromosomes with the following parameters: insert size = 400; standard deviation = 50; read length = 150bp; coverage = 30X.

The resulting dataset contains 586 x 10^6^ reads split into two fastq files. With reads of length 150bp, the index size for the human genome is equal to 120 GBytes.

### B. upVC validation

To better estimate the variant calling quality, we measure the following metrics:

- True Positive (TP): existing variants found by upVC
- False Negative (FN): variants not found by upVC
- False Positive (FP): non existing variants found by upVC

From these values, the following metrics can be computed:

- Precision (P): TP / (TP+FP)
- Sensitivity (S): TP / (TP+FN)

We run both upVC and GATK. The following table summarizes the quality.

GATK has been run with standard parameters. Compared to upVC, the quality is clearly better. However, upVC provides excellent results and legitimates our variant calling implementation on PIM architecture, knowing that the current code has plenty of room for improvement.

### C. Execution time

The experiments have been done on an Intel® Xeon® Silver 4110 CPU @ 2.1 Ghz, 8 cores with 64 GBytes of RAM, equipped with 10 additional UPMEM double-rank DIMM devices with DPU running at 266 MHz. Of a total of 1280 available DPUs (10 x 128), only 1024 full operational DPUs have been used. The available PIM memory size is thus equal to 64 GBytes (64 Mbytes per DPU).

The maximum space allocated to store the index per DPU is about 48 Mbytes, that is 48 GBytes for 1024 DPUs. The human genome index (120 GBytes) requires then 3 passes.

The upVC software has been run with packets of paired-end reads equal to 2^18^ (or 2^19^ = 524288 reads). The average measured execution time for the different tasks of the main loop are the following:

|  | 1024 DPU system |
| --- | --- |
| **Get reads** | 0.32 sec |
| **Dispatch reads** | 0.12 sec |
| **Map reads** | 3.21 sec |
| **Store map results** | 0.18 sec |
| **Update** | 0.53 sec |

The number of loop iterations depends of the number of reads to process and is equal to:

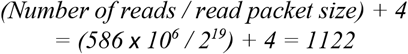

The execution time of one iteration is given by the slowest task, here the mapping (3.21 sec). Consequently, the total execution time of the loop (*T_LOOP_*) is:

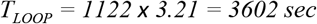

Note that this time must be augmented by the time (*T_INDEX_*) for downloading part of the index before each pass, that is around 40 GBytes. With SSD drives having a bandwidth of 500 Mbytes/sec, 80 seconds must then be added to each pass. The last step of the process is the creation of the VCF file. It’s a very quick operation that takes less than two seconds (*T_VCF_*).

The complete execution time (*T_UPVC_*), that requires 3 passes to process the human genome, is finally given by:

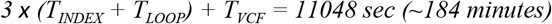

### D. Extrapolation to future 5120 DPU systems

In this section we extrapolate the real results obtained with a 1024 DPU system running at 266 MHz to a 5120 system running at 600 MHz, the target frequency of the next generation of DPUs.

On such a system, that can be seen as a standard bioinformatics server, the full human genome index fits the PIM memory. The whole variant calling process will thus be done in a single pass, leading to the following execution time:

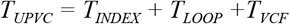

*T_INDEX_* is the time for downloading the 120 GBytes index to the memory DPU (240 sec). The execution time of T_LOOP_ will mainly depend on the execution time of the mapping task since each DPU will store a smaller index (120 GBytes/5120 = 23 MBytes) and then will have proportionally less work to do (23/40 = 0.57). DPU will also run faster (600/266 = 2.25). The mapping task can be approximated by:

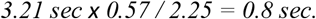

The mapping task still remains the slowest task of the loop, giving a total loop execution time *T_LOOP_* = 898 sec, leading to:

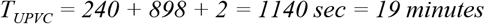

Considering a server dedicated to human variant calling, the index doesn’t need to be reloaded for each new variant calling task. Once downloaded and stored into the DPU memories, the following variant calling tasks will be performed in 15 minutes.

This experimentation clearly indicates that in this parallel upVC implementation, the mapping task performed by the DPUs is by far the longest. The other tasks that run concurrently on the CPU cores are shorter, and lead to an approximate 70% inactivity rate of the host processor. In future implementation, this idle time will be used to consolidate the calling step in order to improve the quality of the results.

## V. Comparison with alternative systems

We compare the performances of the upVC PIM implementation with two other hardware accelerators that efficiently implement the variant calling process: Illumina DRAGEN using proprietary software on 8 FPGA Xilinx UltraScale Plus 16 nm FPGA [13], and Nvidia Parabricks using BWA-GATK4 on 8 NVIDIA®Tesla®V100 GPUs [19].

To provide a fair comparison, we extend performance results to different PIM configurations with increased density and DPU frequency. At the time of writing, the reference platform available at UPMEM is a 2*Xeon Silver 4108 with 128GB RAM and 160 GB of PIM memory with 2560 DPUs clocked at 400MHz. Servers with higher PIM DIMM density such as the Cooper Lake 4* Xeon Gold 6328H with 5120 DPUs and the AMD ARM Epyc with 3584 DPUs are in the process of qualification while DPUs clocked at 500MHz and more are under development.

Performances of these systems are analysed following the three following criteria:

1. Execution time
2. Power consumption
3. Total Cost of Ownership (TCO)

### A. Execution time

Figure 4 reports the execution time of the different systems to process a typical variant calling operation on a 30X human genome dataset. The loading of the reference genome in MRAM is not considered as part of the computation time if enough PIM memory in a system allows single batch runs of upVC. In this case, the reference genome loading process only happens at the start of the server and can be used for all subsequent sample analysis.

**Figure 4:**
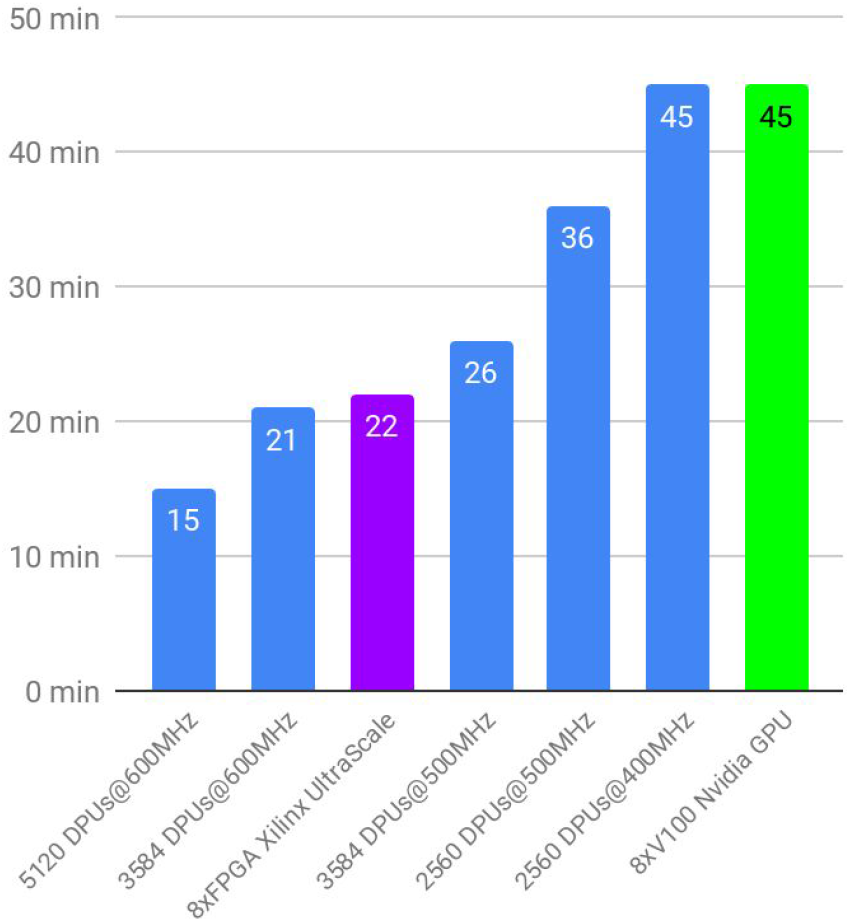
Execution time based on 30X human genome dataset on FPGA, GPU and PIM

A 2560 DPUs configuration does not allow enough space in MRAM to load the entire genome and simultaneously retain enough space to save the reads in MRAM. One DPU has 45,5 MB available to store reference neighbors, which totalizes 116,5 GB with 2560 memory banks, slightly less than the 120 GB of the reference genome. To avoid the need for 2 batches and consequently load 2 halves of the reference genome with long HDD transfers we divide the input read buffer by 2. This way we free enough space for the reference genome but still has for consequence to double the DPU processing time on this configuration. Naturally we observe a gain in performance once the memory space issue is alleviated in 3000+ DPUs configurations.

### B. Power consumption

Figure 5 gives the power consumption of FPGA, GPU and PIM systems.

**Figure 5:**
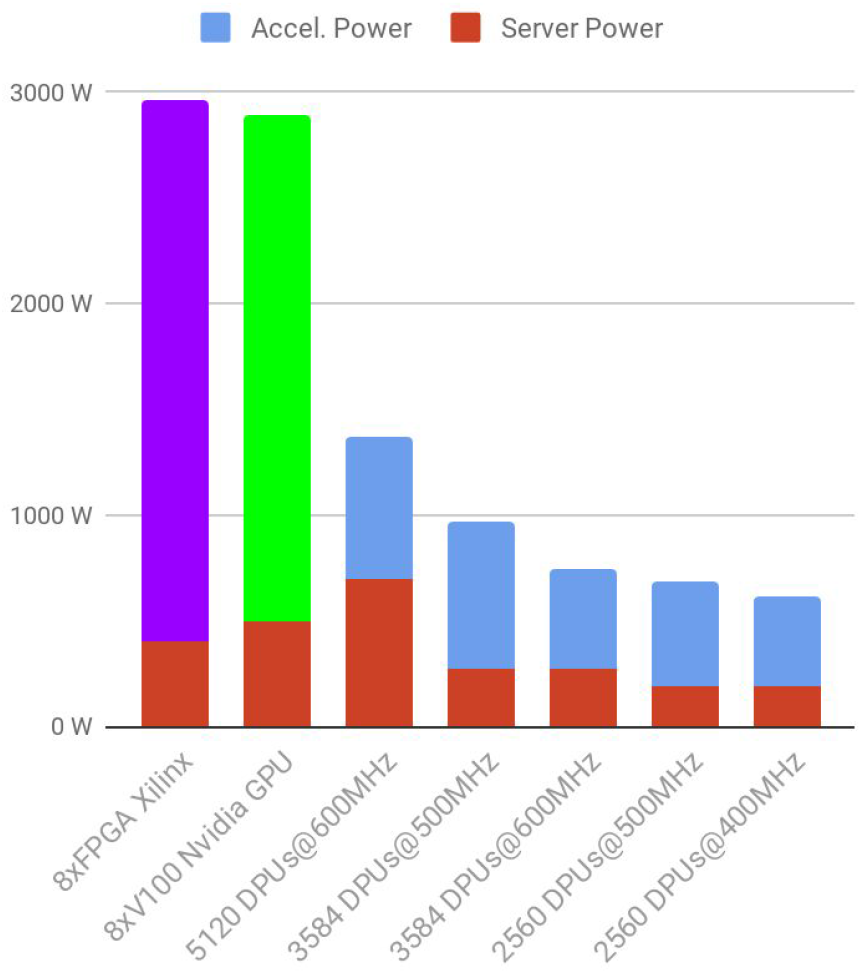
*Power consumption (hardware accelerator + server) of PIM, GPU (8* NVIDIA®Tesla®V100) *and FPGA (8* FPGA Xilinx UltraScale) systems.

The consumption of an FPGA board depends on its configuration. For this workload it is estimated to be used near maximum capacity at 320W per board, 90% of its TDP. The consumption of a V100 GPU is provided by Nvidia and reaches around 300W in full use. UPMEM provides precise measurements of a DPU power consumption and depends on its version and clocking. At 400Mhz, a DPU in current version v1.2 consumes 160 mW, while it consumes 190 mW at 500MHz. DPUs at 600MHz are benefiting from energy reduction designs and are expected at around 120mW. The overall consumption accounts that the charge of DPUs can hardly go over 90% during the entire execution and that every PIM module consumes 3W. PIM based configurations are in average 6x less energy consuming than the considered alternative accelerators.

The consumptions of the server for PIM configurations is based on the 2*Xeon Silver 4110 with 128GB RAM. The TDP is given at 190W. An AMD 2*Epyc has a TDP around 280W and a Cooper Lake with 4* Xeon Gold has a TDP of 700W. We consider a 2*Xeon Gold 6328H server base for alternative accelerators to ensure efficient orchestration at TDP of 400W. 3W per 8 GB of memory accounts for the DRAM consumption.

### C. Total Cost of Ownership (TCO)

Server and infrastructure costs follow the comprehensive AWS TCO cost estimator [24]. The estimator accounts for an annual maintenance of 15% of the hardware cost. The same logic if applied to each of the considered accelerator’s hardware. The consumption evaluation is based on previous energy considerations for a full 3 years and accounts for a cooling and infrastructure overhead (additional 70% of the hardware consumption). The cost of electricity is based on median US commercial price [25]: $0,1/kWh.

**Figure 6:**
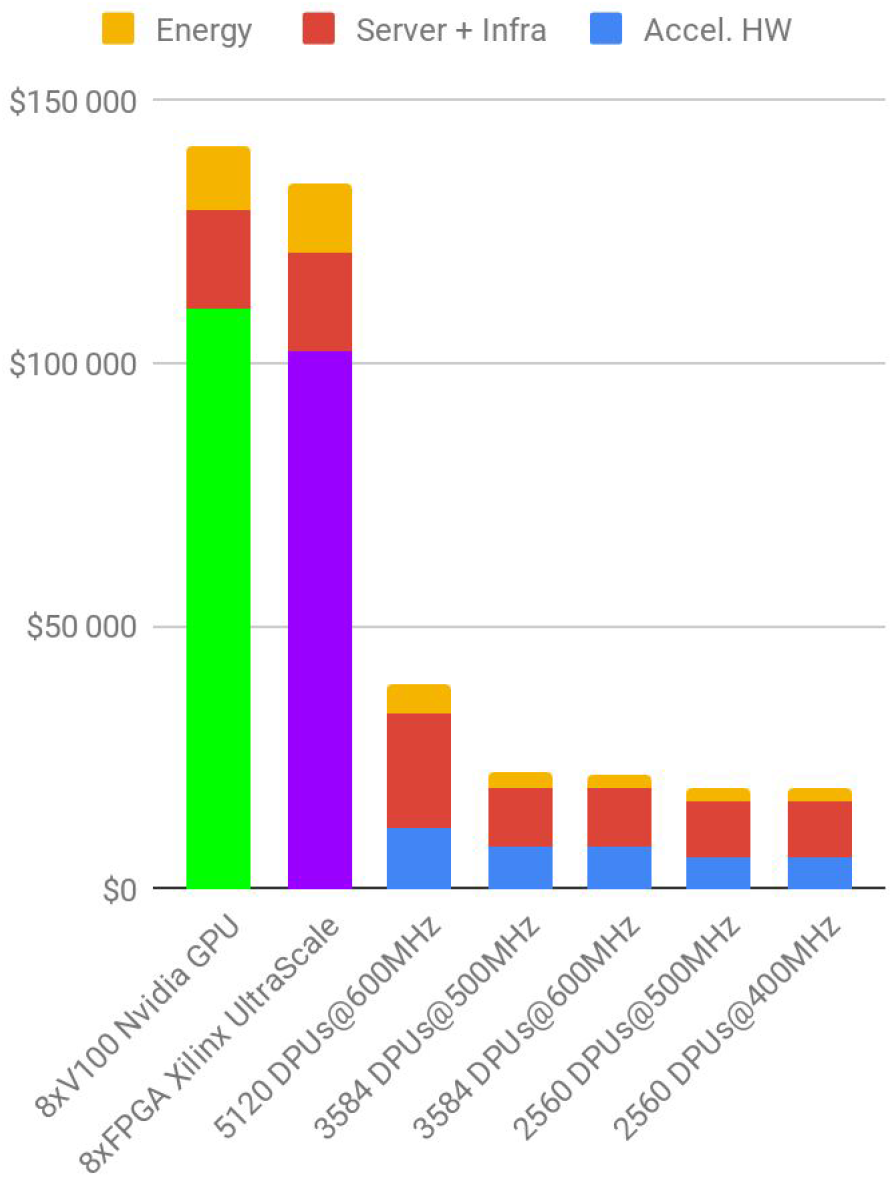
years TCO Comparison between PIM configurations against GPU and FPGA based accelerators for Mapping+ Variant calling.

A full 3 years on premise FPGA based solution’s TCO nears $135,000, which accounts for FPGA board at around $8,800 per unit. It is about 5 times more expensive than UPMEM PIM solution at identical throughput for a 4096 DPUs at 500MHz configuration. Note that if we were to consider the software cost for running Illumina Dragen, an additional $572,000 would be required over a period of 3 years, multiplying the cost reduction made by UPMEM solution by yet again a factor 5.

Nvidia quotes its V100 GPU at $9,500 per unit resulting in a full 3 years TCO of Nvidia Parabrick estimation over $140,000. In terms of algorithms, they are identical to BWA-GATK4, using DNA-Bricks to port them over GPU architectures and do not represent an overhead cost to use the solution. At equivalent throughput it is about 8 times more costly than UPMEM PIM FASTQ to VCF using 4096 DPUs at 500MHz.

Thus, UPMEM technology offers a drastic financial and environmental gain compared to both Nvidia and Illumina solutions. Though it does not reach an as high accuracy, development efforts on upVC are progressively narrowing the gap.

## VI. Conclusion

This implementation demonstrates the performance of the PIM architecture when dedicated to a large scale and highly parallel task in genomics: every DPU independently computes read mapping against his fragment of the reference genome while the variant calling is pipelined on the host.

The algorithm works well within the confines of the experiment but still remains a long way from a real-world application with a holistic alignment strategy. It is a prototype that verifies the capabilities of a PIM architecture in the context of mapping and variant calling. The low CPU usage of this implementation allows additional CPU based functions to complete the pre-variant calling workflow that would pave the road towards a commercial application.

In comparison to existing accelerators, the PIM solution promises to deliver equal to better performances but with massive energy reduction and TCO gains. It is a crucial advantage in sight of the prominent place that genomics is about to occupy in the world of data computing and for its accessibility by medical institutions across the globe. A configuration with 3584 DPUs at 600MHz has the best TCO profile and could bring the cost of human genome analysis near $0,34/genome.

PIM is a promising technology that shows a great potential to solve some of the challenges of genomics in terms of actionable computing power, programmability and cost.

## Footnotes

1 ftp.ncbi.nlm.gov/genomes/H_sapiens/Assembled_chromosomes/seq/

2 ftp.nci.nlm.gov/snp/organisms/human_9606_b150_GRCh38p7/VCF/

